# Genomic and Chemical Diversity of Commercially Available Industrial Hemp Accessions

**DOI:** 10.1101/2021.03.17.435786

**Authors:** Matthew S. Johnson, Jason G. Wallace

**Affiliations:** Institute of Plant Breeding, Genetics, and Genomics, University of Georgia, Athens, Georgia, USA; Department of Crop & Soil Sciences, University of Georgia, Athens, Georgia, USA

## Abstract

High consumer demand for cannabidiol (CBD) has made industrial hemp (*Cannabis sativa*) an extremely high-value crop. However, high demand has resulted in the industry developing faster than the research, resulting in the sale of many hemp accessions with inconsistent performance and chemical profiles. To determine the genetic and phenotypic consistency in available CBD hemp varieties, we obtained seed or clones from 22 different named accessions. Genotypes (~48,000 SNPs) and chemical profiles (% CBD and THC by dry weight) were determined for up to 8 plants per accession. Many accessions--including several with the same name--showed little consistency either genetically or chemically. Most seed-grown accessions deviated significantly from their purported levels of CBD and THC based on the supplied certificates of analysis. Several also showed evidence of an active tetrahydrocannabinolic acid (THCa) synthase gene, leading to unacceptably high levels of THC in female flowers. We conclude that the current market for CBD-hemp varieties is highly unreliable, making many purchases risky for growers. We suggest options for addressing these issues, such using unique names and developing seed and plant certification programs to ensure the availability of high-quality, verified planting materials.

## Introduction

Hemp (*Cannabis sativa)* is a dioecious annual plant that is believed to have been domesticated around 6000 years ago in China, with some evidence of use as far back as 12,000 years ago ^1–3^. Different hemp varieties have been used for fiber, seeds, medicine, and recreation for thousands of years ^4^, and hemp has recently been used to produce biofuels ^5^, plastics ^6,7^, and building composites ^8,9^.

Similar to other crops, different hemp varieties serve specific uses. However, until recently the United States had banned all hemp varieties from commercial production due some of them being used as recreational drugs (marijuana) ^10^. The psychoactive properties of marijuana are due to high amounts of a specific secondary metabolite, tetrahydrocannabinol (THC). Recognizing that not all hemp is created equal, the 2018 US Farm Bill allowed growers to cultivate “industrial” hemp (defined as varieties with <0.3% THC by dry weight) throughout the US ^11^. This has led to a surge of interest in hemp production, especially for varieties with high production of other, non-psychoactive metabolites (cannabinoids). Currently, the largest interest is in varieties bred to produce cannabidiol (CBD), a non-psychoactive cannabinoid used as a medicine and health food supplement. Both CBD and THC derive from the same precursor, cannabigerolic acid (CBGa)^12,13^. They are most concentrated in the trichomes of female flowers, as are the over 100 other known cannabinoids that are present at much lower concentrations^14,15^.

The most well-studied application of CBD is to control seizures, which is the basis of the FDA-approved drug epidiolex. CBD is also marketed as a nutritional supplement to help with anxiety, pain, depression, and sleep; most of these claims are anecdotal, although there are some studies supporting its usefulness ^16–18^. Regardless of efficacy, the market value of CBD products is currently estimated at over $4.7 billion dollars per year ^19^.

### Hemp genetics

*C. sativa* has a diploid genome (2n=20) with an estimated size of 818 Mb for female plants and 843 Mb for males ^20^. Sex determination in hemp is via an imperfect XY genetic system, where plants with an XX chromosome pair are generally female and ones with XY are generally male. However, environmental conditions, plant hormones, and ethylene inhibitors can affect the sexual phenotype of the plants, resulting in genetically female plants with at least some male flowers ^21–26^. The rate at which genetically female plants produce male flowers is controlled by many genetic factors and is still not well understood ^27,28^, though some grain varieties have been bred to produce both male and female flowers on one plant to maximize seed production ^29^. Pollination lowers cannabinoid yield by ~50% ^30^, however, so growers interested in these compounds use “feminized” seed or clones of female plants. (Feminized seed is produced from two genetically female plants, one of which has been chemically treated to produce male flowers) ^23,31^. Even with these precautions some plants may still produce male flowers; these plants are killed when found so as to minimize pollination.

Since its legal status prevented most research on hemp until recently, it is practically an orphan crop, with few genomic resources and almost no public germplasm collections. Despite its lack of resources, there have been significant advances in understanding hemp genetics for several key traits, including cannabinoid production ^32–35^, sex expression ^36^, fiber quality ^37^, and population structure and diversity ^38–40^. Most recently, multiple draft reference and draft genomes have become available ^41,42^, bringing hemp into the genomics era.

### Issues with commercial hemp cultivation

Although interest in commercial hemp cultivation has exploded since the 2018 Farm Bill, many issues of naming and quality control plague the field. Because all varieties of hemp were outlawed for several decades, breeding and naming of varieties has been largely clandestine and ad-hoc, with names frequently recycled to reflect the most successful or desirable cultivars. Thus there is no guarantee that the variety “Cherry Wine” received from one supplier is the same as--or even related to--a variety of the same name from a different supplier. In addition, standards for CBD and THC production are lacking. This is particularly important because any plants with >0.3% THC are classified as marijuana and must be destroyed, potentially costing farmers significant lost revenue. Some companies provide certificates of authenticity (COAs) that attest to how much of each compound a variety will produce. (See the Discussion for issues related to these COAs.)

To determine the extent of these issues in the current hemp market, we surveyed the genetic and chemical diversity of twenty-two commercially available hemp accessions. We identified both the genetic relationships among the accessions and the genetic consistency within an accession. We also tested the production of CBD and THC for each line and compared these to industry and legal standards, along with the provided Certificates of Authenticity. These comparisons are not meant to evaluate specific sources or accessions *per se*, but rather to demonstrate the overall state of the market and give an idea of how reliable (or not) it is for interested growers.

## Materials and Methods

### Plant material

Twenty-two commercial hemp accessions were purchased or donated from various sources (Table 1). Since accessions frequently pass among groups, it is impossible to say if these companies are the original sources, or if they have done any selections to differentiate them from the original source. This collection focused on cannabidiol production, but accessions for fiber and seed were also included. Twenty of the accessions were distributed as seeds (some feminized, others not) and two of them were clonally propagated. For our experiments, all clonal plants were propagated from a single original plant to ensure that each replicate was an exact genetic copy.

**Table 1:**
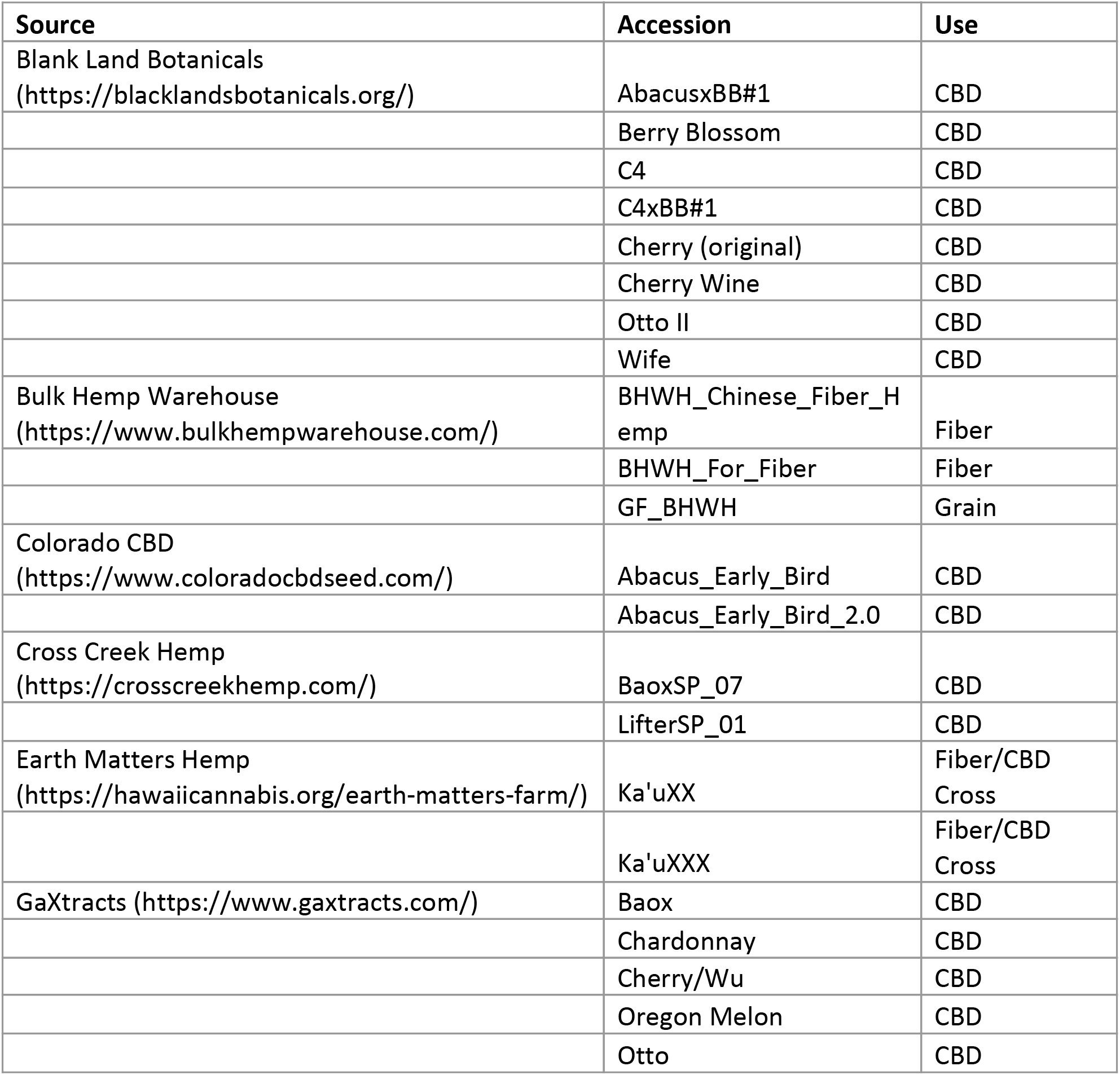
Accessions used in this study

Seeds were soaked in water for 12 hours to initiate germination. Due to low germination rates, 15 seeds were planted per half-gallon pot and thinned to only 1 plant per pot starting at 2 weeks. Clones were made by cutting a seven-inch section of stem from the mother plant, trimming off all leaves and growing points except the topmost one, and dipping in cloneX (Growth Technology) rooting solutions before planting into half-gallon pots. All plants were grown in a commercial potting medium (Sun Gro **Metro Mix 830**).

Plants were fertilized twice a week using a diluted 20-20-20 fertilizer (1000 ppms) and a diluted micronutrient mixture (Jackpot Micronutrient Mixture; 500 ppm). To maintain the plants in a controlled vegetative state, growth conditions were kept under an 18-hour light / 6-hour dark cycle. All plants were grown in greenhouses at the University of Georgia (Athens, Georgia).

### Genomic Data

Ten leaf punches were taken from each plant and sent to LGC Genomics for DNA extraction and genotyping-by-sequencing (GBS) ^43^ with restriction enzyme MslI. Paired-end 150bp reads were generated using Illumina NextSeq V500/550. Libraries were demultiplexed using the Illumina bcl2fastq software (version 2.17.14). SNPs were aligned with BWA mem ^44^ and SNPs called with BCFtools ^45^ requiring a minimum base quality of 20 and only outputting SNPs (not indels). All bioinformatic scripts (including exact parameters used) are available at https://github.com/wallacelab/paper-johnson-hemp-gs, and adaptor- and restriction-fragment-verified sequencing data is available at NCBI under Bioproject PRJNA707556.

Raw SNPs were then filtered in a series of steps. Misalignments and low-coverage sites were filtered out by removing all sites with an average genotyping depth of <15 reads per individual, and paralogs were removed by filtering out sites with >125 average depth (based on initial data exploration). We then removed sites with minor allele frequencies <2.5% (since most of these are sequencing errors) and with >10% heterozygosity (since these are often paralogs being misaligned to the same location). This resulted in 48,029 SNPs in the final dataset. The phylogenetic tree was created from these SNPs using the neighbor-joining method in TASSEL v5.2.40 ^46^.

### Cannabinoid analysis

Fifty-two days after sowing, eight replicates of each accession were placed into a flower room with a 12-hour light / 12-hour dark cycle to initiate flowering. Plants were laid out in a randomized complete block design. Any plants that showed male flowers were removed from the room to eliminate pollination. The remaining female plants were kept in the flowering room for 12 weeks, at which point the panicles in the top six inches of each plant were harvested, trimmed of excess leaf material, placed in a paper bag, and dried at 35°C for 2 weeks.

Approximately 200 mg of dried flower material was weighed out for each sample and the exact weight recorded. Samples were placed in 20 ml of methanol and agitated for 3 minutes to extract the cannabinoids. 1 ml of the supernatant was passed through a 0.22 um filter, and 50 μl of filtrate was diluted into 950 μl of methanol, resulting in a 400x dilution of the original samples.

These diluted samples were assayed by high-performance liquid chromatography (HPLC) on a Shimadzu LC2030C HPLC machine. The components used for the HPLC analysis were the NexLeaf CBX column (2.7 µm, 4.6 × 150mm; part number: 220-91525-70), NexLeaf CBX guard column (part number: 220-91525-72), eleven-cannabinoid standard mix (part number: 220-91239-21), and high sensitivity method solvents A (0.085% phosphoric acid in water) and B (0.085% phosphoric acid in acetonitrile) (part number: 220-91394-81). The flow rate was 1.5 ml/min with a gradient starting from 30% solvent/70% solvent B and ramping to 5% solvent A/95% solvent B over 8 minutes. Injection volume was 5μL, and a guard column temperature of 35° Celsius was maintained by an internal oven. Standard curves were generated for each target cannabinoid with minimum correlation coefficients (R^2^) of 0.999 over the six concentration levels (0.5ppm, 1ppm, 5ppm, 10ppm, 50ppm, and 100ppm). The original sample weights were used to determine the precise cannabinoid concentration in the original sample.

### Genomic Prediction

Genomic prediction was performed using basic GBLUP ^47^ analysis in TASSEL ^46^. Accuracy was assessed with 1000 iterations of 10-fold cross-validation.

### Figures

All figures were made in R using packages argparse v2.0.1 ^48^, ggplot2 v3.3.0 ^49^, gridExtra v2.3 ^50^, and tidyverse ^51^.

## Results

We planted 8 replicates of the 22 accessions, resulting in 176 total pots. Of these, 3 had no seeds germinate despite having 15 seeds originally planted, and 25 developed male flowers and so were removed from the experiment. (The males were from 11 different accessions, none of which claimed to be feminized seed.)

All 148 remaining female plants were genotyped with Genotyping-By-Sequencing (GBS), resulting in >48,000 markers after filtering (see Methods). Phylogenetic clustering showed some expected patterns, such as the fiber accessions clustering together and clones clustering tightly to each other. (Figure 1A). Minor differences among clones are presumably due to a low level of sequencing errors that made it through our filters.

**Figure 1:**
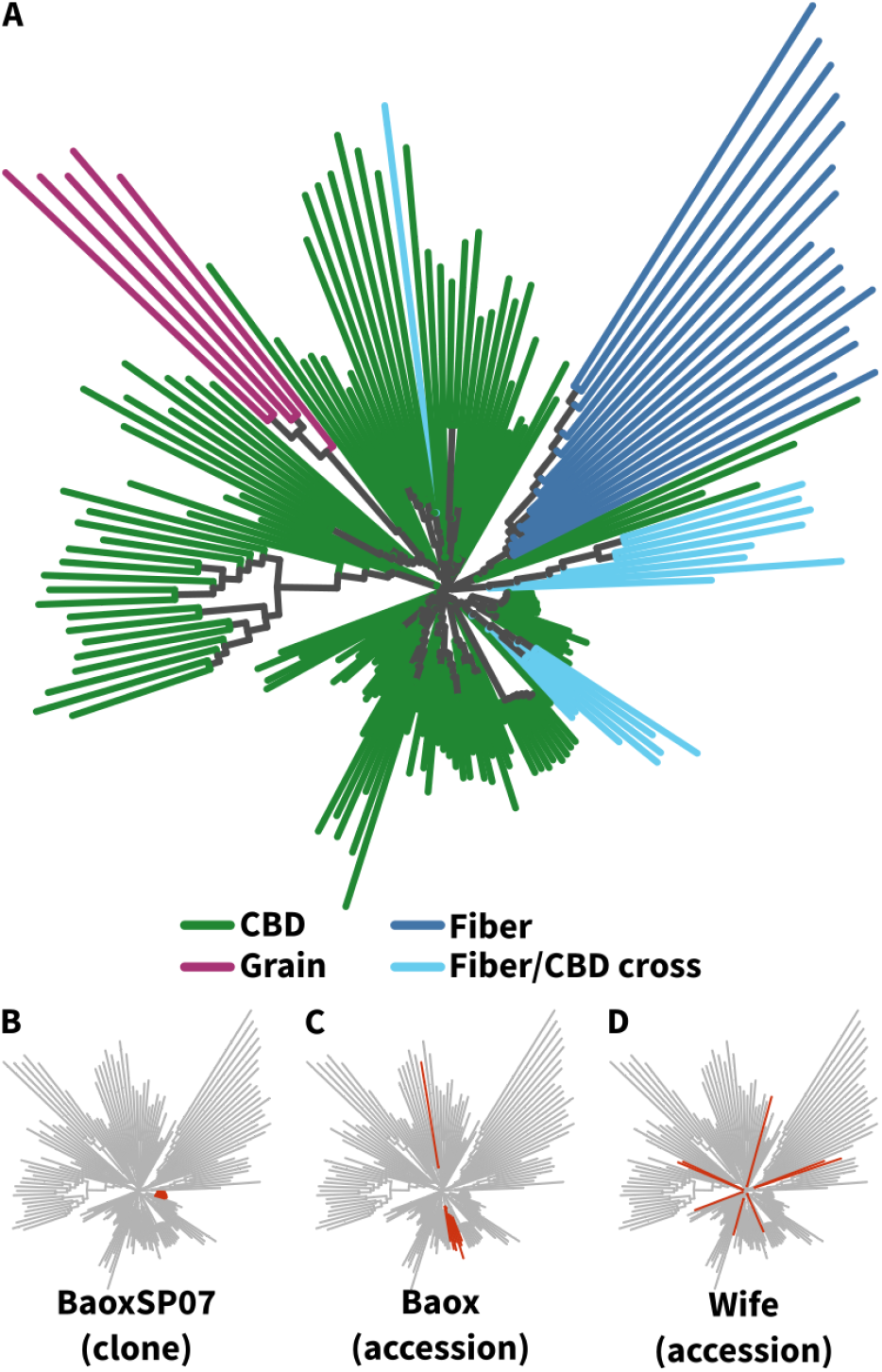
Relationships among accessions based on a neighbor-joining tree of SNP data. (A) All plants (173 plants from 22 accessions). Branch tips are colored by use type. (B-D) Variation among individual accessions. All individuals within the clonal variety “BaoxSP07” (B) cluster tightly, whereas the seed accession “Baox” shows more variation, including one off-type. Importantly, although both B and C are supposedly Baox, they do not cluster with each other on the tree. (D) The “Wife” accession showed extreme variation, with individual plants scattered across the entire tree. (See Supplemental Figure S1 for more details on individual accessions.)

Most of the seed-grown CBD accessions showed little consistency. For example, two accessions named “Baox” (“Baox” and “BaoxSP07,” from 2 different suppliers) showed no real relationship to each other, and even within one accession individual plants spread out among the tree (Figure 1B; Supplemental Figure S1).

Of the 148 female plants, eleven had flowers that did not properly develop, leaving 137 flower samples tested for cannabinoid content. THC levels ranged from undetectable up to 11.08% THC by dry weight. Eighty-nine plants produced flowers with more than the legal limit of 0.3% THC (Figure 2A), including 15 plants with >1% THC and one with >10% THC (Figure 2A). CBD levels ranged from undetectable (mostly fiber varieties) up to 16.7% dry weight (Figure 2B). Twelve plants produced THC without any CBD, including three of them with >1% THC by dry weight, though all of these were fiber varieties.

**Figure 2:**
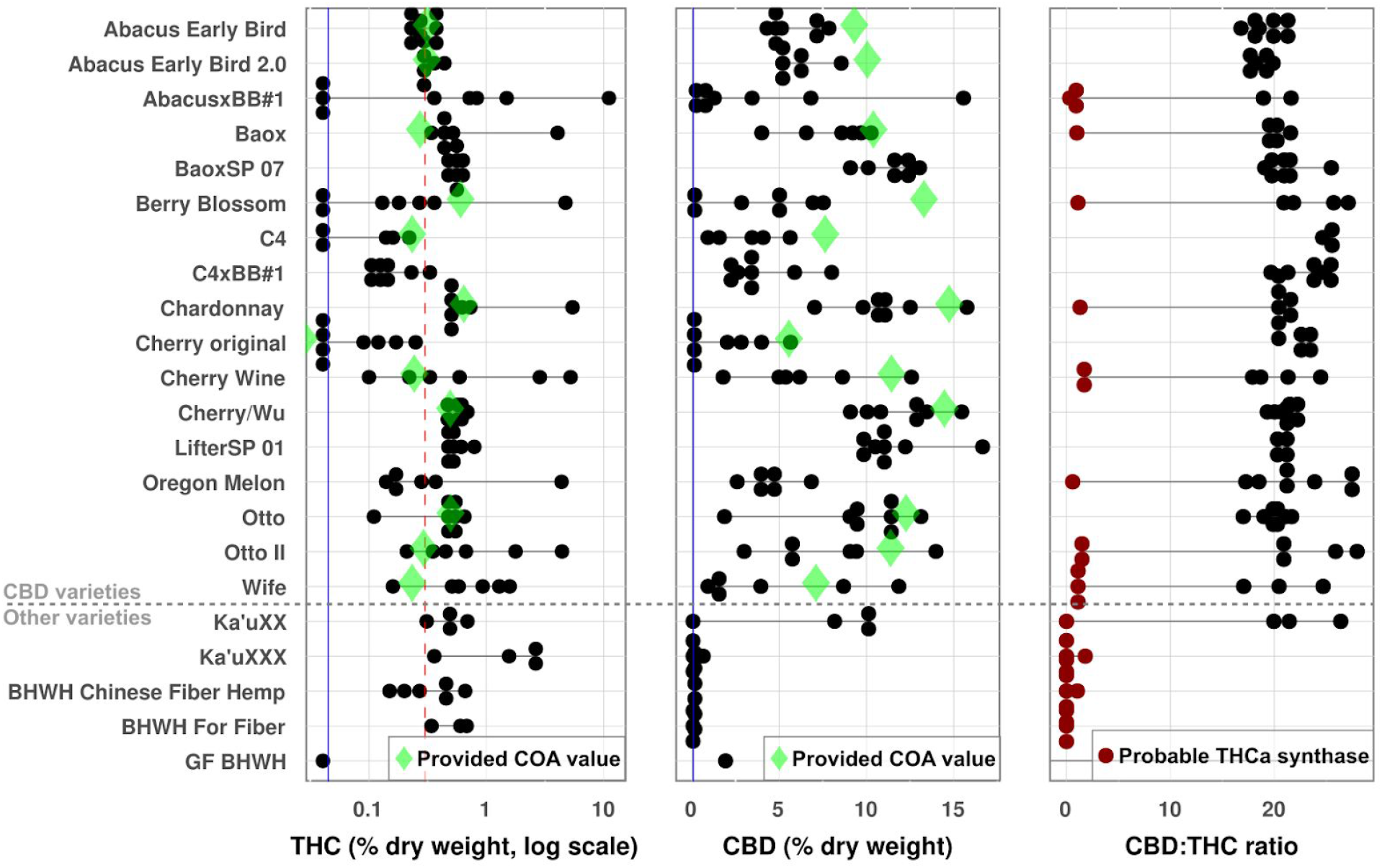
Cannabinoid Analysis. THC and CBD were determined for all plants as a percent dry weight of harvested material. Each dot represents a single plant, and horizontal black lines show the range of values for a given accession. Green diamonds show the value provided on the Certificate of Analysis (COA) for each variety, and vertical blue lines show the limit of detection. (A) THC values. The red dotted line is maximum allowable value in the United States (0.3%); plants above this line would be considered marijuana. (B) CBD. (C) CBD:THC ratio. Red dots indicate plants whose CBD:THC ratio indicates the presence of at least one active THCa synthase gene.

The CBD:THC ratios varied from ~0 (for the plants that produced no CBD) up to ~28:1 (Figure 2C). Some plants produced no detectable THC (and thus have no ratio), but these plants also had very low levels of CBD (Supplemental Table S1). The CBD:THC ratio can indicate which genes are present in the plant, with ratios of ~20:1 indicating no active THCa synthase and ratios of ~2:1 indicating one active THCa synthase gene ^28^. Based on this, twenty-eight plants had ratios that indicate the potential presence of a copy of the THCa synthase gene (Figure 2C).

Twelve accessions came with a certificate of analysis (COA), which suppliers use to show what level of cannabinoid production should be expected from the plant. These certificates are important for growers to know that their crop will remain under the 0.3% legal limit for THC, along with estimating the return on investment for CBD. However, most accessions had less CBD than their COA showed, and almost all of them had more THC (Figure 2), both of which could potentially cause issues for commercial growers.

With an eye toward breeding applications, we used both the genetic and phenotypic data to perform genomic prediction among the plants for CBD, THC, and their ratio. Prediction accuracies (assessed by 10-fold cross-validation) were universally low, however, with average accuracy of only 3-7% (Supplemental Figure S2; Supplemental Table S2).

## Discussion

The current federal regulations have created a fine line between legal hemp and illegal marijuana. Inconsistencies in plant genetics can greatly complicate the already complex process of legally growing hemp. Although planting clones ensures the highest consistency, many farmers choose to plant seeds because of their much lower cost. This lower cost comes with risk due to inconsistent plant genetics and seed feminization that can make it difficult to produce hemp profitably ^30^.

### Seed Feminization

Many farmers have received seeds that were improperly feminized or not feminized at all, resulting in lost revenue and lawsuits ^52^. All feminized product used in this experiment produced only female flowers, although the numbers were too small to draw definite conclusions from this. (Farmers consider a seed lot to be well feminized if less than 1 in 4000 plants produce male flowers; personal observation.)

### Genetic Relationships

Most accessions tested showed little genetic consistency (Figure 1; Supplemental Figure S1), which likely explains their phenotypic inconsistency (Figure 2). As expected, clonal accessions were the major exception, though some seed accessions (like Chardonnay) showed good within-accession consistency. This indicates that at least some accessions from some suppliers are reliable, although without extensive testing it is impossible to say which. Conversely, plant accessions with the same name but from different suppliers did not show any genetic clustering (Figure 2B), meaning that they are actually no more related than any two random accessions. Growers should be careful when purchasing materials; the best approach is probably to just assume that each supplier is selling completely different seed regardless of what they name it.

### THC production

The most concerning results from this experiment were the number of plants that produced excessively high levels of THC. The CBDa synthase gene naturally produces low levels of THC ^53^, so any plant producing CBD will have some amount of THC. However, plants with as much or more THC production than CBD almost certainly have an active THCa synthase gene ^28^. Eight CBD accessions had at least one plant that showed evidence of an active THCa synthase gene (Figure 2), even though all were supposed to be low-THC varieties. The presence of an active THCa synthase gene in CBD-production lines is very concerning, and the rate was surprisingly high (28 of 121 seed-grown plants). More extensive testing would be needed to see if any of the other CBD accessions also contain plants with active THCa synthase. All four fiber accessions also showed evidence of active THCa synthase, though in this case the issue is less one of CBD production and more their possible contamination of other seed lots. Farmers may want to invest some time and resources into testing small batches of seeds from different suppliers to identify which ones are the most stable and trustworthy (not to mention well performing).

### CBDa Synthase Produces THC

As previously mentioned, CBDa synthase naturally produces low levels of THC, which explains why almost all the plants tested showed some level of THC (Supplemental Table S1). Even with THC much below CBD production, the plants which produced the highest levels of CBD all exceeded the federal level of 0.3% THC at full maturity. This implies that, with current varieties, there may be a limit to how much CBD a plant can produce while staying below the legal limit of THC. In the long run this limit might be improved by using natural or induced variation in the CBDa synthase gene to select for more specific enzyme variants. For now, however, frequent testing of plants as the flowers mature can help farmers see when their plants are getting too close to that limit and adjust their harvest times accordingly.

### Certificates of Authenticity

One concerning pattern we noticed was that several COAs were printed so that they show misleadingly low levels of THC. Specifically, they highlighted the low levels of Δ9-THC (the actual psychoactive form) while de-emphasizing THCa (the acid precursor that is decarboxylated into Δ9-THC by heat). US federal testing guidelines require including both^54,55^, and using the official formula of [total THC] = [Δ9-THC] + 0.877 * [THCa]^55^, only 7 of the 14 accessions with COAs were actually compliant with US regulations. (This is a separate issue from our results in Figure 2 differing from the COAs, since that could be due to differences in growing conditions.) Some of these COAs may have been issued before the interim final rule that established these guidelines (31 Oct 2019)^54^, and we would hope they have been updated since. Nonetheless, growers should pay close attention when ordering materials and ensure that the product information is reported accurately.

### Genomic Prediction

Our attempts at genomic prediction did not yield useful models. This is likely due to the phenotypic variability present, since in theory we have the exact genotype of every plant tested. Given the great potential for genomic selection and other methods (such as Speed Breeding ^56,57^) to accelerate hemp breeding, further work should be done to improve phenotype quality and prediction accuracy. Larger training populations, using clones to exactly replicate each plant (even those originally from seed), deeper genotyping coverage, and other adjustments could be done to improve both the precision and accuracy of genomic prediction on these materials. A specific benefit of implementing genomic selection in hemp would be the ability to reliably select male lines, since it is not possible to directly phenotype them. (This is actually analogous to dairy cattle breeding, where genomic selection was first pioneered to select (male) bulls for improved (female) milk production ^58^.)

## Conclusion

The hemp industry is experiencing many growing pains associated with its rapid development in the last six years. There are issues with genetic stability, economic viability, and governmental regulations. Despite these issues, the market continues to grow year after year, and interest in this crop continues to expand. With the support that this crop receives from consumers and the support it is beginning to receive from a wide range of researchers, there is a great opportunity for hemp to play an increasingly important role in a wide range of industries. Given the variability we saw both among and within accessions, some sort of standardization is needed so that producers can be confident in the material they receive. A good first step is for suppliers to start using unique names for each of their accessions. Not only will this help clarify the market, it will also allow each company to capitalize on branding their own unique varieties. A more complex but badly needed step is an industry seed-certification process. Certified seeds will allow growers to purchase with confidence. Some states are already moving forward with their own seed certifications ^59,60^, but until rigorous, independent verification is implemented across the industry, growers face the prospect of getting a bad lot any time they purchase from a new supplier. Ultimately, these and other changes will need to occur to make the market for hemp varieties robust and reliable enough for the long-term.

## Acknowledgments

We thank Tim Coolong for assistance in obtaining hemp material. This work was supported by the UGA Institute of Plant Breeding, Genetics, and Genomics, and the UGA Cultivar Development Program.

