## Supplemental Figure for "Genomic and Chemical Diversity of Commercially Available Industrial Hemp Accessions"

### Supplemental Data

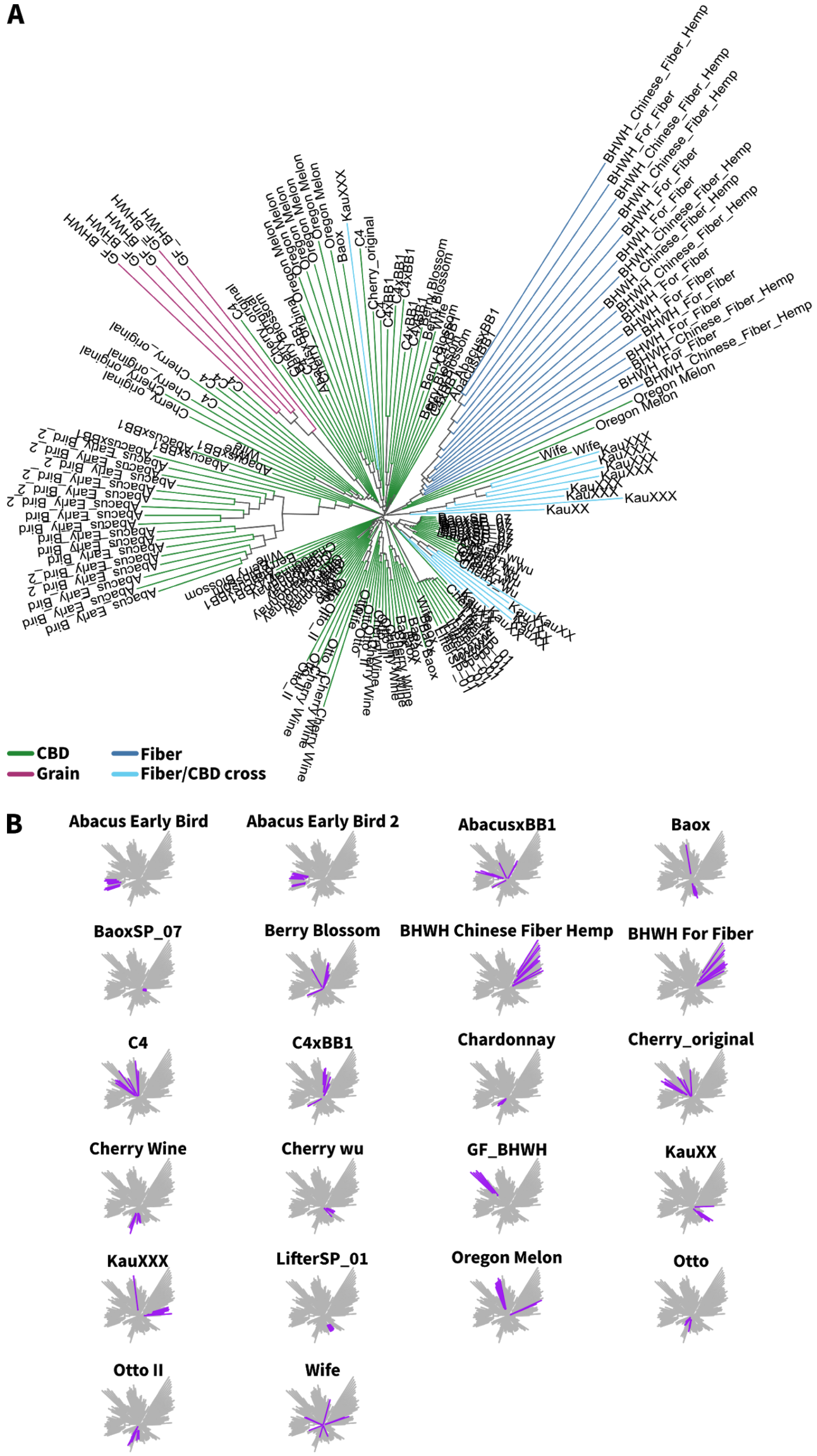


**Supplemental Figure S1 - Phylogenetic trees.** (A) Neighbor-joining tree of all genotyped plants with labeled tips. (B) The same tree as in (A), but with each accession highlighted. Some accessions are tightly clustered, while others are scattered across the tree.

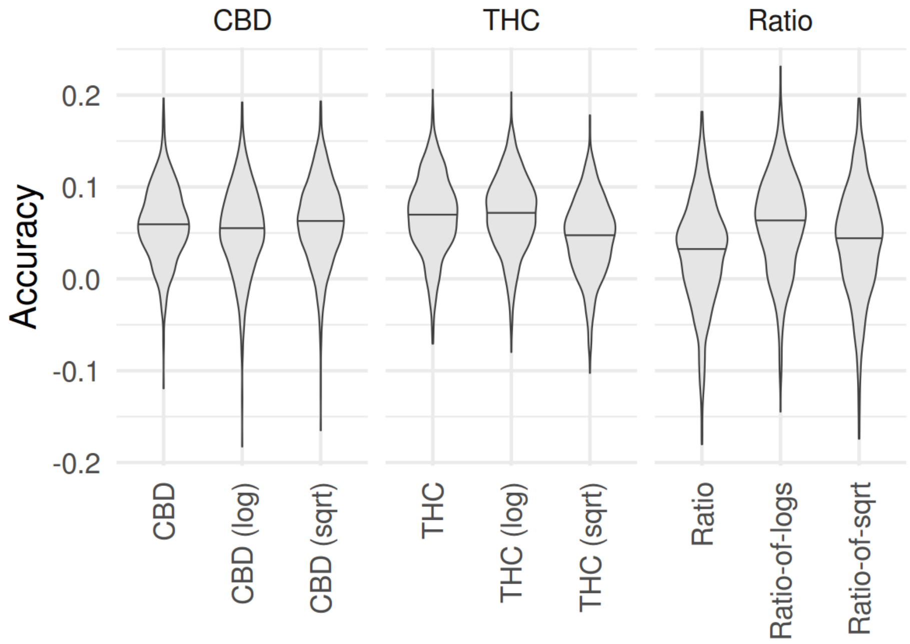


**Supplemental Figure S2 - Genomic Prediction Accuracy.** GBLUP genomic prediction was performed for CBD, THC, and their ratio across 148 plants. We tested the raw phenotype data and log- and square root (sqrt)-transformed data, but in all cases accuracy was low (<0.1). Distributions show the range of average prediction accuracy across 1000 ten-fold cross-validation runs.

**Supplemental Table S2: Genomic Prediction Accuracy from 1000 Ten-fold Cross-Validations**

| Trait | Mean Accuracy ± St. Dev. |
| --- | --- |
| CBD | 0.06±0.04 |
| CBD_log | 0.054±0.047 |
| CBD_sqrt | 0.062±0.045 |
| THC | 0.069±0.043 |
| THC_log | 0.07±0.041 |
| THC_sqrt | 0.046±0.042 |
| Ratio | 0.029±0.058 |
| Ratio_of_logs | 0.06±0.053 |
| Ratio_of_sqrt | 0.041±0.059 |
